# The novel roles of bovine milk-derived exosomes on skin anti-aging

**DOI:** 10.1101/2023.03.23.532505

**Authors:** Lu Lu, Wei Bai, Miao Wang, Chunle Han, Huanqing Du, Na Wang, Mengya Gao, Dan Li, Fengwei Dong, Xiaohu Ge

## Abstract

Exosomes are small vesicles released from cells and present in various mammal biological fluids, such as bovine milk, which worked for skin care for many years besides dairy. In addition, Exosomes were regarded as a vehicle for intercellular communication. Therefore, we aimed to investigate the novel roles of bovine milk-derived exosomes (MK-Exo) on human skin anti-aging. Purified MK-Exo can be directly uptake by the keratinocytes and fibroblast *in vitro* and upregulate the expression of the natural factors related to skin moisturizing, including Filaggrin (FLG), Aquaporin 3 (AQP3), CD44 in the keratinocytes and hyaluronidase (HAS2) in the fibroblast, and MK-Exo promoted the cell migration of the fibroblast, while rescue its expression of type I collagen (Col I), type III collagen (Col III) after ultraviolet radiation. Furthermore, the phototoxicity test, photoallergy test, repeated skin irritation test, skin allergy test, and patch test confirm the safety of MK-Exo on the skin. Finally, the roles of MK-Exo in preserving moisture and anti-wrinkle were also identified in humans. Then, MK-Exo was smeared on the facial skin of 31 female volunteers twice a day for 28 days, and the functions were evaluated following the safety assessment *in vivo*. These studies reveal the novel roles of bovine milk-derived exosomes in human skin aging, which opens a new way of skin care.

## Introduction

Bovine milk is known to be used as a raw material in the food industry. It is also widely used in cosmetic industries due to its considerable biological potential, mainly derived from casein and whey proteins [1, 2]. Milk-based products positively affect skin conditions, including improved wound healing, elasticity, and moisturizing when topically applied in creams, and ointments, et al. [3–5].

Skin is a protective layer of the body of any animal, including humans. As age progresses, specific changes occur in the skin, the most visible signs of which include wrinkles, dryness, and loss of natural smoothness [6, 7]. Although skin aging is a complex biological process, the mechanism is not yet completely understood; it is commonly considered that intrinsic factors of free radical toxicity, hormonal reduction, mitochondrial DNA damage, extrinsic factors of UV, and lifestyle are responsible for skin aging [8–11]. Several anti-aging strategies are developed, such as skin care, moisturizing preparation, and anti-wrinkling treatment. Wrinkles are mainly caused by a lack of elastic features of the skin, so inhibition of elastic fiber degradation is used for anti-wrinkling [12]. Hyaluronic acid and sericin were also used to preserve the skin’s hydration for anti-wrinkling [13, 14].

Exosomes (40–150 nm in diameter) are one subtype of extracellular vesicle originating in the endocytotic compartment and released via biological membrane fusion between the multi-vesicular bodies and the cell membranes from nearly all kinds of cells. They contain proteins, lipids, nucleic acids, and other biological molecules and work as intercellular communication tools to regulate the properties of target cells [15–17]. The released exosomes widely exist in intercellular space and various bodily fluids, including bovine milk [18].

Some studies have shown that exosomes distributed in the skin also play a role in skin conditions through the intercellular crosstalk of various skin cells. For example, Hu et al. found that human dermal fibroblast-derived exosomes could ameliorate skin photoaging [19]. Liu et al. discovered that exosomes derived from keratinocytes could regulate melanocyte pigmentation via loaded microRNA [20]. Besides exosomes originally from the skin, Kim et al. found that ectopic exosomes derived from milk can suppress melanogenesis [21].

Bovine milk is rich in exosomes. Although the application of bovine milk-derived ingredients is widely accepted as functional cosmetics, there still needs to be more studies on the roles of bovine milk-derived exosomes in skin conditions.

In this study, we found that bovine milk-derived exosomes(MK-Exo) can up-regulate the expression of FLG and CD44 in keratinocytes and HAS2 in fibroblasts and arrest the UV-triggered collagen reduction. Meanwhile, bovine milk-derived exosomes can also significantly improve the cell migration of fibroblasts. Followed by the skin toxicity tests in animals and humans, we further validated the functions of bovine milk-derived exosomes moisturizing and anti-wrinkling in female volunteers. These results indicate that the bovine milk-derived exosomes are safe and may work as a novel ingredient for anti-aging skin.

## Results

### Characterization and labeling of the exosomes isolated from bovine milk

Owing to abundant non-EV proteins, sugars, milk fat, and other components, whole bovine milk is a highly complex material for isolating pure MK-Exo. Therefore, we combined acid precipitation and density gradient ultracentrifugation to isolate MK-Exo (Figure. 1A). We chose fresh bovine milk to avoid any unexpected risk from industrial food processing. Multiple steps of centrifugation clarified the supernatant of acid precipitation. We employed morphology and protein markers (CD9 and TSG101) to identify the MK-Exo existing fractions (Figure. 1B, C) following DC. The MK-Exo collection was further identified by size distribution and exosomal positive and negative markers (Figure. 1D, F). The slight peak of HPLC indicates the high purity of MK-Exo (Figure. 1G) in the collections.

**Fig. 1.**
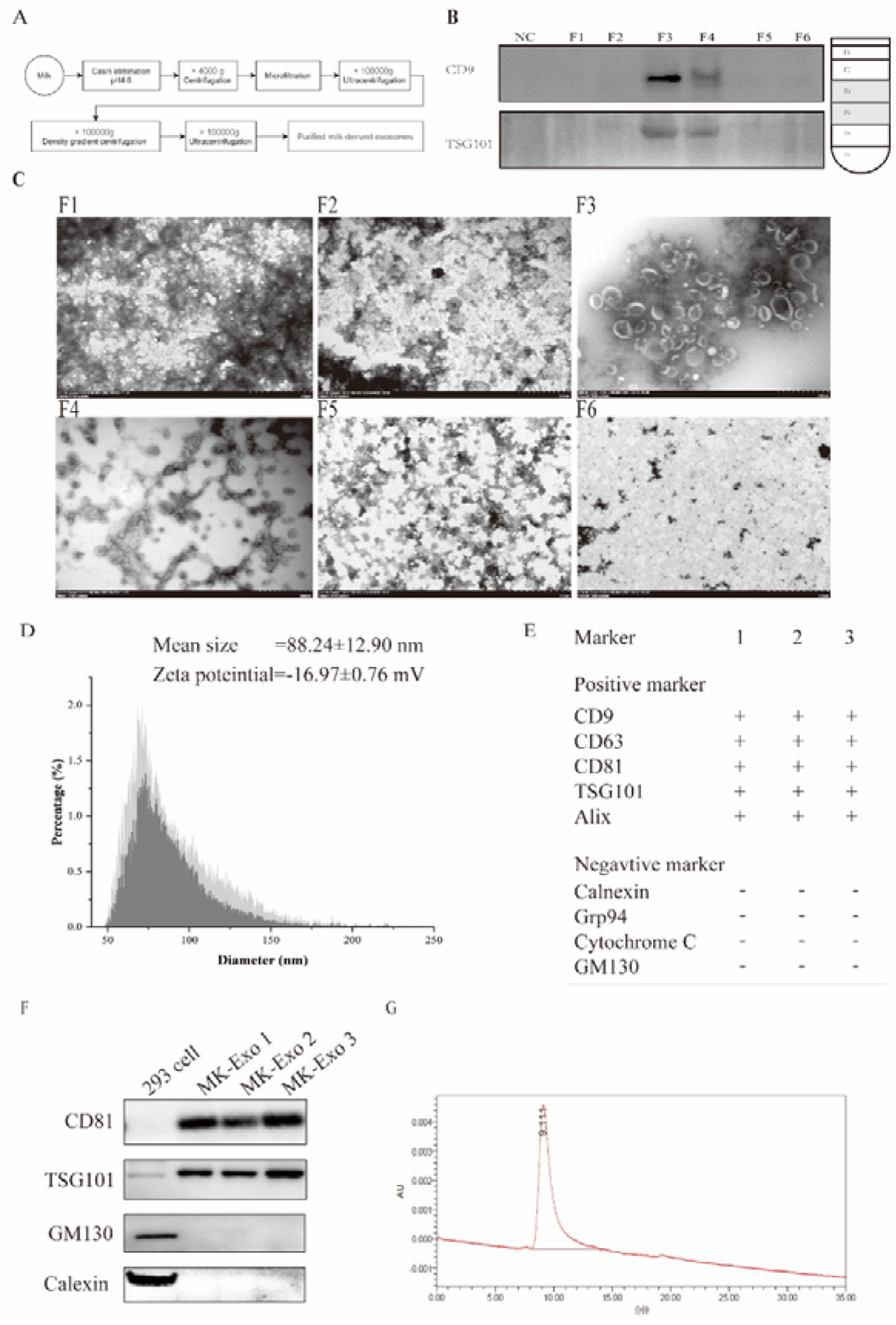
Preparation and characterization of MK-Exo. (A) Flow chart of the preparation of MK-Exo by density gradient ultracentrifugation. Each fraction of density gradient ultracentrifugation was analyzed by Western blot of CD9, TSG101 (B), and the morphological characteristics of exosomes in MK-Exo were identified by TEM (C). (D) Mixed the fraction of F3 and F4, particle size was determined by NanoFCM. (E) LC-MS analyzed positive markers and negative markers of exosomes. (F) The expression of CD81, TSG101, GM130, and Calnexin was analyzed by Western blot. (G) HPLC identified the purity of MK-Exo.

The hydrophobic fluorescent dye is widely used in labeling exosomes for tracing *in vitro* and *in vivo*. Still, we found that the labeling efficiency of exosomes with different dyes is highly varied, from 15.7% to more than 90% (Supplementary Figure. 1A, D). Therefore, to accurately trace the exosomes, we labeled MK-Exo using AIE, which has the highest efficiency for further study.

### The effects of MK-Exo on human keratinocytes

Keratinocytes are constructional skin cells and take part in moisturizing [22]. To investigate whether MK-Exo can influence the moisturizing functions of the keratinocytes, we first incubated the MK-Exo with the HaCat cells. The cells could up-take the MK-Exo (Figure. 2A). The preliminary result indicated that MK-Exo might function across the species on the human keratinocytes.

**Fig. 2.**
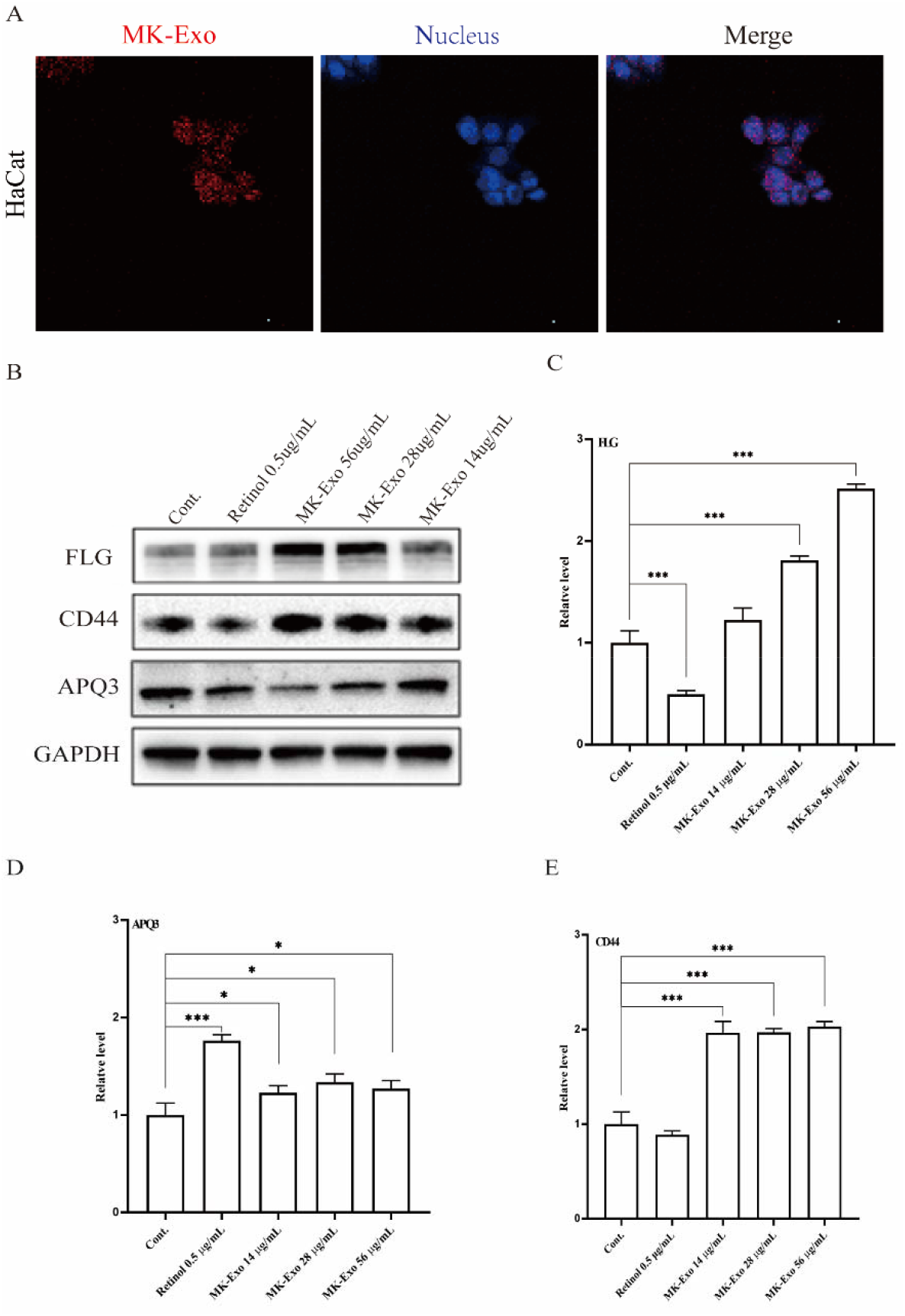
The effect of MK-Exo on HaCat. (A) Representative fluorescence images of AIE-labeled MK-Exo uptake by HaCaT. (B) The protein expression levels of FLG APQ3 and CD44 were determined by Western blot. The gene of FLG (C), APQ3 (D), and CD44 (E) transcript levels were determined by RT-qPCR. Asterisk (*) represented P < 0.05; a double asterisk (**) represented P < 0.01; a triple asterisk (***) represented P < 0.001.

To further investigate whether MK-Exo could trigger gene expression change related to moisture in keratinocytes *in vitro*, we explored the mRNA and protein levels of a few relevant genes considered as the indicator for moisturizing. We found that MK-Exo was able to elevate the status of filaggrin (FLG), a natural moisturizer, up to around three times (Figure. 2B, C), and CD44, the receptor of HA, up to more than 60% (Figure. 2B, E). Aquaporin 3 (AQP3) is the most abundant skin aquaporin that facilitates water and glycerin transport into SC to help keep it hydrated. However, we found that MK-Exo reduced the level of AQP3 at relatively high concentrations (Figure. 2B, D). The results indicate that MK-Exo may work as moisture by inducing the expression of FLG and CD44 in keratinocytes without a change in AQP3 expression.

### The effects of MK-Exo on human fibroblasts

Fibroblasts are constructional skin cells and participate in skin conditions, including moisturizing and anti-wrinkling. To investigate whether MK-Exo can influence the functions of the fibroblasts, we first incubated the MK-Exo with the CCC-ESF-1 cells. We found that the cells also could up-take the MK-Exo (Figure. 3A). The preliminary result suggested that MK-Exo also might function across the species on the human skin fibroblasts.

**Fig. 3.**
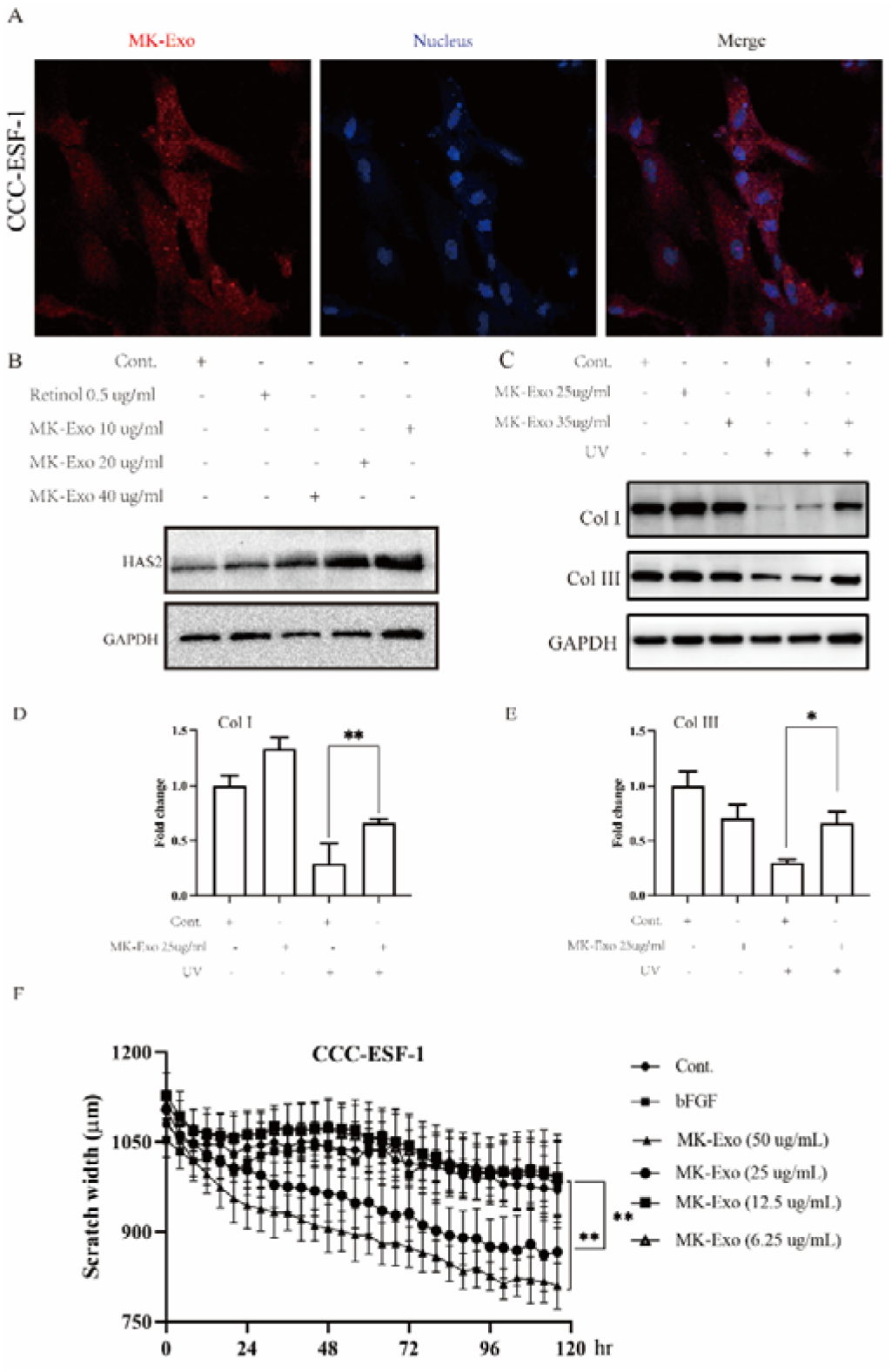
The effect of MK-Exo on CCC-ESF-1. (A) Representative fluorescence images of AIE-labeled MK-Exo uptake by CCC-ESF-1. (B) The protein expression levels of HAS2 and GAPDH were analyzed by Western blot. The expression of Col I and Col III were determined by Western blot (C) and RT-qPCR (D-E). (F) The varies of scratch width at different time points. (Asterisk (*) represented P < 0.05; a double asterisk (**) meant P < 0.01.)

To further investigate whether MK-Exo could influence the moisturizing skin function of human fibroblasts *in vitro*, we explored the protein levels of hyaluronan synthases (HAS2). We found that MK-Exo could elevate the level of HAS2 more than two times (Figure. 3B). More HAS2 indicates that more hyaluronan acid will be synthesized. The result suggests that MK-Exo also could play the role of moisturizing by inducing the production of hyaluronan acid, which is also one kind of natural moisture factor, by HAS2.

Skin aging is primarily due to alterations in the dermal extracellular matrix, significantly decreasing collagen I content. Meanwhile, about 80% of facial aging is attributed to ultraviolet radiation exposure from sunlight, which also could introduce collagen degradation [23]. However, MK-Exo did not significantly change the expression level of collagen I and III; instead, it can moderate collagen I and III reductions in fibroblasts following UV exposure (Figure. 3C, E). In addition, we found that MK-Exo also could improve fibroblast cell migration (Figure. 3F, Supplementary Figure. 2) instead of keratinocytes (Data not shown). These results suggested that MK-Exo may play the role of anti-wrinkling by reducing collagen degradation and improving cell migration.

### Primary safety evaluation of MK-Exo on skin

As the potential to be used as a cosmetic material, skin contact is one of the most common routes to MK-Exo during their intended daily use. Nevertheless, the toxic effects of cutaneous MK-Exo exposure on animal and human skin remain unexplored. Therefore, first, we preliminary evaluated the toxicity of MK-Exo on the skin in animals by combing skin allergy tests, skin photoallergy tests, repeated skin irritation tests, and skin photo-irritation tests. No allergic reaction occurred following the sensitization by MK-Exo in the skin allergy tests, skin photoallergy tests (Table. 1), and repeated skin irritation tests (Supplementary Table. 2). Meanwhile, no skin damage occurred on the skin in the skin photo-irritation tests (Supplementary Table. 3).

**Table 1.** Skin allerge test and skin photoallergy test

| Group | Time (h) | Erythema |  |  |  |  | Edema |  |  |  | Quantity<br>(Score ≥ 2) | Sensitization<br>rate (%) |
| --- | --- | --- | --- | --- | --- | --- | --- | --- | --- | --- | --- | --- |
|  |  | 0 | 1 | 2 | 3 | 4 | 0 | 1 | 2 | 3 |  |  |
| Skin allergic test |  |  |  |  |  |  |  |  |  |  |  |  |
| NC | 24 | 10/10 | 0 | 0 | 0 | 0 | 10/10 | 0 | 0 | 0 | 0 | 0 |
|  | 48 | 10/10 | 0 | 0 | 0 | 0 | 10/10 | 0 | 0 | 0 | 0 | 0 |
| PC | 24 | 0 | 9/20 | 11/20 | 0 | 0 | 0 | 8/20 | 12/20 | 0 | 20 | 100 |
|  | 48 | 2/20 | 13/20 | 5/20 | 0 | 0 | 1/20 | 13/20 | 6/20 | 0 | 17 | 85 |
| MK-Exo | 24 | 20/20 | 0 | 0 | 0 | 0 | 0 | 0 | 0 | 0 | 0 | 0 |
|  | 48 | 20/20 | 0 | 0 | 0 | 0 | 0 | 0 | 0 | 0 | 0 | 0 |
| Skin photoallergy test |  |  |  |  |  |  |  |  |  |  |  |  |
| NC | 24 | 10/10 | 0 | 0 | 0 | 0 | 10/10 | 0 | 0 | 0 |  | 0 |
|  | 48 | 10/10 | 0 | 0 | 0 | 0 | 10/10 | 0 | 0 | 0 |  | 0 |
| PC | 24 | 0 | 7/10 | 3/10 | 0 | 0 | 2/10 | 8/10 | 0 | 0 |  | 80 |
|  | 48 | 2/10 | 8/10 | 0 | 0 | 0 | 6/10 | 4/10 | 0 | 0 |  | 40 |
| MK-Exo | 24 | 10/10 | 0 | 0 | 0 | 0 | 10/10 | 0 | 0 | 0 |  | 0 |
|  | 48 | 10/10 | 0 | 0 | 0 | 0 | 10/10 | 0 | 0 | 0 |  | 0 |

To further confirm the potential sensitization on human skin, we recruit 31 female volunteers to evaluate MK-Exo sensitization by a patch test. None of the skin reactions were observed at 0.5hr, 24hr, and 48hr (Supplementary Table. 4) in all the volunteers.

Overall, the results indicated the absence of sensitization and irritant potential of MK-Exo on animal and human skin, and there will be no risk in efficacy evaluation in humans.

The proportion of responding animals to the number of tested animals in the erythema and edema column of the skin reaction score (0, 1, 2, 3, 4), and the sensitization rate is the percentage of the number of animals with a response score of 2 or more to the total number of animals in that group.

### The anti-aging effect of MK-Exo in human

To confirm the potential effect of MK-Exo in anti-aging through moisturizing and anti-wrinkle in humans, we recruited 31 female volunteers aged 26 to 45 years old. After the patch tests, everyone facial applied the MK-Exo twice daily for 28 days without using other cosmetics. The skin conditions were detected on day 2, day 14, and day 28 separately. The skin’s moisture content increased by 4.64% on day 14 and 5.6% on day 28 in all the cohorts. Interestingly, the increase of moisture content is higher in the volunteers aged 36-45, while the increase in volunteers aged 26-35 is not statistically significant. (Table 2). The skin brightness also dramatically increased on day 28. More than 90% satisfy the moisturizing effect of MK-Exo (Data not shown). In addition, the photo of the volunteer’s facial skin gloss increased on day 28. The results indicated that MK-Exo has the function of moisturizing.

**Table 2.** Skin moisturizing test and skin wrinkle test

|  | Age | D2 | D14 | D28 |
| --- | --- | --- | --- | --- |
| Moisture content | 26-45 (n=31) | +1.66% | +4.64%* | +5.60%* |
|  | 26-35 (n=14) | +3.23% | +4.04% | +4.66% |
|  | 36-45 (n=17) | +1.39% | +7.16%* | +7.96%* |
| F3/F4 | 26-45 (n=31) | - | +5.35%* | +6.33%* |
|  | 26-35 (n=14) | - | +6.15% | +3.66% |
|  | 36-45 (n=17) | - | +6.29%* | +10.74%** |
| R2 | 26-45 (n=31) | - | +2.76% | +7.24%** |
|  | 26-35 (n=14) | - | +0.12% | +2.62% |
|  | 36-45 (n=17) | - | +5.37% | +11.80%** |
| Amount of wrinkles | 26-45 (n=31) | - | -5.27%*** | -4.99%*** |
|  | 26-35 (n=14) | - | -7.24%** | -6.77%** |
|  | 36-45 (n=17) | - | -3.65%*** | -3.52%** |
| Wrinkle area | 26-45 (n=31) | - | -9.37%*** | -9.59%*** |
|  | 26-35 (n=14) | - | -7.67%** | -10.01%** |
|  | 36-45 (n=17) | - | -10.59%*** | -9.29%*** |

(Asterisk (\*) represented $P < 0.05$ ; a double asterisk (\*\*) meant $P < 0.01$ ; a triple asterisk (\*\*\*) represented $P < 0.001$ .)

Skin elasticity was also detected on day 14 and day 28. We found that F3/F4 value and R2 value increased by 6.33% and 7.24% on day 28, with a higher increase of 10.74% in the cohort of age 36-45 (Table. 2). In addition, the wrinkle area and amount of wrinkles were detected and reduced by 9.37% and 5.27% on day 14 and by 9.59% and 4.99 % on day 28. The results suggested that MK-Exo could also play the role of anti-wrinkle in humans.

All results indicated that MK-Exo was safe and may moisturize and anti-wrinkle by improving the expression of FLG, CD44 in keratinocytes, and HAS2 in fibroblasts, enhancing cell migration and inhibiting the UV-induced reduction of collagen in fibroblasts.

## Discussion

It’s the first study to combine and evaluate the safety and efficacy of MK-Exo in the skins of animals and humans and the first evidence supporting that MK-Exo could participate in skin care by anti-aging. It is also the first discovery that MK-Exo could change gene expressions such as FLG, CD44, HAS2, and collagen. This study may provide shreds of more solid evidence for the potential of MK-Exo as cosmetic material. However, it is not yet clear how MK-Exo works and what is the critical molecular player. MK-Exo consists of thousands of types of proteins and non-coding RNAs. Omics technology can be employed to explore the key players [24–26], the specific proteins or RNAs which induce the changes in the given biological processes of the skin, in further studies. The key players will be the critical quality attributes and stand at the center of the quality control system in the industrial process.

Interestingly, we found the MK-Exo function differently on cohorts of different ages in some test items. MK-Exo may have a stronger position on skin anti-aging in older females. Indeed, more subjects may be needed for confirmation. However, the difference might suggest that different age cohorts have specific skin conditions and respond differently to the MK-Exo. The mechanism should be explored deeply.

In addition, exosomes are a novel potential non-virus delivery system for small molecular compounds, peptides, proteins, and nuclear acids in the pharmaceutical industry [27–30]. Because of unlimited supply, low cost, and simple oral route, MK-Exo may be one optimal candidate for the drug delivery system. However, it is generally believed that large-scale production and quality control standards may hinder industry progress. Fortunately, we have erased the central problems in the production process. In addition to drugs, MK-Exo can also work as a delivery system in the cosmetic industry. Crossing the skin barrier is a critical issue for most functional cosmetic ingredients. Meanwhile, MK-Exo function in human skin indicates that it can cross the skin barrier, at least partially. Therefore, MK-Exo could also work as a delivery system for other available cosmetic materials crossing the skin and improving their efficacy.

## Materials and Methods

### 2.1 Exosomes preparation

Fresh bovine milk was obtained from a local dairy factory. Exosomes were isolated by density gradient centrifugation. Briefly, the pH of milk was adjusted to pH 4.6 by hydrochloric acid (Merck) followed by centrifugation at 4000×rpm (Beckman) at 4□ for 30 min. Then, Sucrose density gradient centrifugation was performed as described previously[31], and the isolated MK-Exo were sterile filtered through a 0.22 μm filter and stored at −80 °C until further use.

### 2.2 Quantitative assessment of protein concentration

The protein concentration of MK-Exo was measured by BCA Kit (Thermo, Waltham Mass, USA) according to the manufacturer’s instructions.

### 2.3 Transmission electron microscope

The MK-Exo (100 μg/mL) was fixed in 2% (w/v) paraformaldehyde at room temperature for 15 min. The mixture (10 uL) was mounted on a formvar-carbon coated grid (Beijing XXBR Technology, Beijing, China) at room temperature for 3 min and then stained by uranyl oxalate solution (4% uranyl acetate, 0.0075 M oxalic acid, pH 7) for 1 min. After the samples were observed by TEM (Hitachi High-Technologies Corporation, Tokyo, Japan).

### 2.4 Size distribution and particle number

The MK-Exo size distribution and number were determined by NanoFCM (NanoFCM Inc., Xiamen, China) according to the user manual.

### 2.5 Western blotting

Western blot analyses were performed according to the previously described protocols [32]. The primary antibodies used in this study were as follows: CD81 (Abcam), TSG101 (BD, 612696), GM130 (Abcam), Calnexin (Abcam), FLG(Santa Cruz), CD44(Thermo), AQP3(Abcam), Col I(Abcam), Col III(Abcam), HAS2(Santa Cruz), GAPDH (Abcam). Depending on the primary antibody, the secondary antibodies were either goat anti-rabbit (Abbexa) or goat anti-mouse antibodies (Protientech). The immunoreactive protein was used with ECL (Thermo) to image immunoblots.

### 2.6 Purity analysis

The sample was analyzed using an SEC-1000 column (7.8*150 mm, 7 um, Thermo). The column was eluted at a 0.3 mL/min flow rate with 150 mM NaCl and 20 mM phosphate buffer pH 7.2.

### 2.7 Proteomic analysis

To identify the proteomics of MK-Exo, liquid chromatography/mass spectroscopy (LC-MS/MS, Thermo) analysis was performed as described before with modification [33]. The LC-MS/MS using Easy NLC 1200-Q Exactive Orbitrap mass spectrometers (Thermo). The nano-HPLC system was equipped with an Acclaim PepMap nano-trap column (C18, 100 Å, 75 μm × 2 cm) and an Acclaim Pepmap RSLC analytical column (C18, 100 Å, 75 μm × 25 cm). One μL of the peptide mix was typically loaded onto the enrichment (trap) column. All spectra were collected in positive mode using full-scan MS spectra scanning in the FT mode from m/z 300-1650 at resolutions of 70000. For MSMS, the 15 most intense ions with charge states ≥2 were isolated with an isolation window of 1.6 m/z and fragmented by HCD with a normalized collision energy of 28. A dynamic exclusion of 30 seconds was applied.

The raw files were searched using Proteome Discover (version 2.4, Thermo) with Sequest as the search engine. Fragment and peptide mass tolerances were set at 20 mDa and 10 ppm, respectively, allowing a maximum of 2 missed cleavage sites. The false discovery rates of proteins and peptides were 1%. DAVID Bioinformatics Resource 2021 (http://david.abcc.ncifcrf.gov/)2 analyzed the differential expression proteins with recommended analytical parameters to identify the most significantly enriched signal transduction pathways in the data set.

### 2.8 Cell lines

The cells of CCC-ESF-1 and HaCaT were purchased From the National Institute of Cell Resource, Beijing, China. CCC-ESF-1 was cultured in DMEM (Gibco, CA, USA) supplemented with 10% FBS (Gibco), 100 μg/mL streptomycin. HaCaT were cultured in MEM (Gibco) supplemented with 10% FBS, 100 μg/mL streptomycin. Cells were maintained in a humidified incubator at 37 □ under an atmosphere of 5% CO2.

### 2.9 RNA isolation

According to the manufacturer’s instructions, total RNA was isolated from cells using Trizol (Thermo).

### 2.10 RT-qPCR

RT-qPCR was performed using PrimeScript™ RT Master Mix (Takara) and SYBR Green Premix Pro Taq HS (Takara). The primers used in this study were listed in the supplementary table 1.

### 2.11 MK-Exo up-taking

2 × 10^4^ cells/well were seeded onto a chamber side and incubated overnight. Then, the cells were treated with AIE labeled MK-Exo (10 μg/mL) for 4 h. after fixation with 4% paraformaldehyde for 20 min. The nucleus was labeled with DAPI (Thermo) and then observed by a confocal laser-scanning microscope.

### 2.12 Cell migration assays

The wound healing assay detected cell migration. Coculture experiments were performed by seeding CCC-ESF-1 (2 × 10^4^ cells/well) and culturing them with different concentrations of MK-Exo for 24 h; the migration rate was measured by quantifying the wound width automatically (CBM, Biotek).

### 2.13 Skin allergy test

The male Dunkin Hartley was randomized into three groups: NC (PBS, n=10), PC (2, 4-dinitrochlorobenzene, n=20), and MK-Exo (n=20), and were topically treated with sample for six h induction at D0, D7 and D14 (NC: 0.2g PBS, PC: 0.2mL (10mg/mL) 2, 4-dinitrochlorobenzene, MK-Exo: 0.2g). Then, on day 28, the sample was applied to the untreated abdomen and the rib area for excitation ( NC: 0.2g MK-Exo, PC: 0.2mL (5mg/mL) 2, 4-dinitrochlorobenzene, MK-Exo: 0.2g), and observed skin responses at different time points (24h, 48h).

### 2.14 Skin Photoallergy test

The male Dunkin Hartley was randomized into three groups: NC (PBS, n=10), PC (2, 4-Methylcoumarin, n=10), and MK-Exo (n=10). First, the light induction phase involved Dunkin Hartley injecting 0.1 mL sensitizer (Freund’s Adjuvant Complete: Normal saline = 1:1) under the four corners of the neck hair removal area and applying samples in the hair removal area (NC: 0.1g PBS, PC: 0.1mL (10%) 2, 4-Methylcoumarin, MK-Exo: 0.1g), after 30 min, UVA (10.2J/cm2) was irradiated once daily for five times. Secondly, Dunkin Hartley divided the two sides of the spine into four acting sites (left 1, 3 and right 2, 4). Next, 3 and 4 were applied samples for 30 min (NC: 0.02g PBS, PC: 0.02 mL (10%) 2, 4-Methylcoumarin, MK-Exo: 0.02g). After that, necks 1 and 3 were covered with tin foil, and UVA irradiation was performed at 10.2 J/cm^2^. Finally, the skin reaction was observed at 24h and 48h.

### 2.15 Skin moisturizing and wrinkle test

The healthy Chinese subjects of female (n=31, age: 26-45) and the MK-Exo was diluted with water to 60 ug/mL and applied morning and evening for 28 days. The efficacy of MK-Exo was measured by VISIA using skin hydration (Corneometer CM825), skin elasticity (Cutometer MPA580), skin wrinkle number, and area (PRIMOS CR).

### 2.16 Statistical analyses

The statistical methods used for each experiment are described in the relevant figure legends. Experiments were performed with at least three replicates, and results were considered statistically significant at p < 0.05.

## Supporting information

Supplementary Mathers

## Acknowledgments

We thank Professor Dan Ding of Nankai University for the gift of AIE. We also thank Ganggang Zhao, Jianxin Yin, Ning Chen, Few li, Zhijun Wen, and Quan Zhang for their excellent technical assistance.

## Funding

Funding was provided by Tingo Exosomes Technology Co., Ltd, Tianjin, China.

## Author contributions

All authors reviewed the manuscript; X.H.G. funding acquisition, conceptualization, supervision; F.W.D. funding acquisition; L.L. conceptualization, validation, formal analysis, writing - review & editing, project administration; W.B. and M.W. methodology, resources, investigation, data curation, visualization, writing an original draft; C.L.H. investigation, formal analysis, data curation, visualization; M.Y.G., and N. W. investigation, data curation; H.Q.D. Project administration, investigation, D.L. visualization, writing an original draft. All authors contributed to the article and approved the submitted version.

## Conflict of interest

The authors declare that the research was conducted in without any commercial or financial relationships that could be construed as a potential conflict of interest.

## Data and materials availability

All data are in the main text or the supplementary materials. Further inquiries can be directed to the corresponding author.

## Ethics statement

The animal study was reviewed and approved by the Experimental Animal Ethics Committee of the CAST (Tianjin) Inspection and Test Co., Ltd. (2021010401).

The research on volunteers was in line with the basic principles of the international declaration of Helsinki. Therefore, it was approved by the local Institutional Review Board and Ethical Committee of the Centre Testing International (Hangzhou) Co., LTD (project identification code: A22200825411010102CR1).

## Notes

### Competing Interest Statement

The authors have declared no competing interest.

