## Supplementary Mathers for "The novel roles of bovine milk-derived exosomes on skin anti-aging"

**Supplementary Materials and Methods**

**Fluorescent labeling**

The MK-Exo were labeled with DID ( Sigma-Aldrich, St. Louis, USA), DIO (Sigma-Aldrich), PKH67 (Sigma-Aldrich), and aggregation-induced emission (AIE) according to the manufacturer’s instructions. The labeling ratio was determined by nanoFCM. AIE was provided as a gift by Prof. Ding Dan Nankai University.

**Repeated skin irritation test**

The female New Zealand rabbits (n=4) were topically treated with 0.5g sample (PBS, MK-Exo) on both sides of the spine for 14 days, respectively; erythema and edema were observed and recorded one h after every time.

**Skin Photoirrritation test**

The Dunkin Hartley (male: n=3, female: n=3) divided the two sides of the spine into four acting sites (left 1, 3 and right 2, 4). 1 and 2 were applied samples for 30 min. After that, 1 and 3 were covered with tin foil, and UVA irradiation was performed at 1000 mj/cm^2^. Finally, skin reactions were observed at different time points (1h, 24h, 48h, 72h).

**Skin Patch test**

The patch test equipment with an area of no more than 50 mm^2^ and a depth of about 1 mm, and add the sample into the patch tester with a dosage of about 0.02 mL - 0.025 mL (NC: PBS, MK-Exo group: MK-Exo). The patch tester was attached to the curved side of the subject's forearm for 24 h. After removing the patch tester and observed the skin reactions at 0.5h, 24h, and 48h.
