## Supplementary figure for "The novel roles of bovine milk-derived exosomes on skin anti-aging"

**Supplementary Figures**

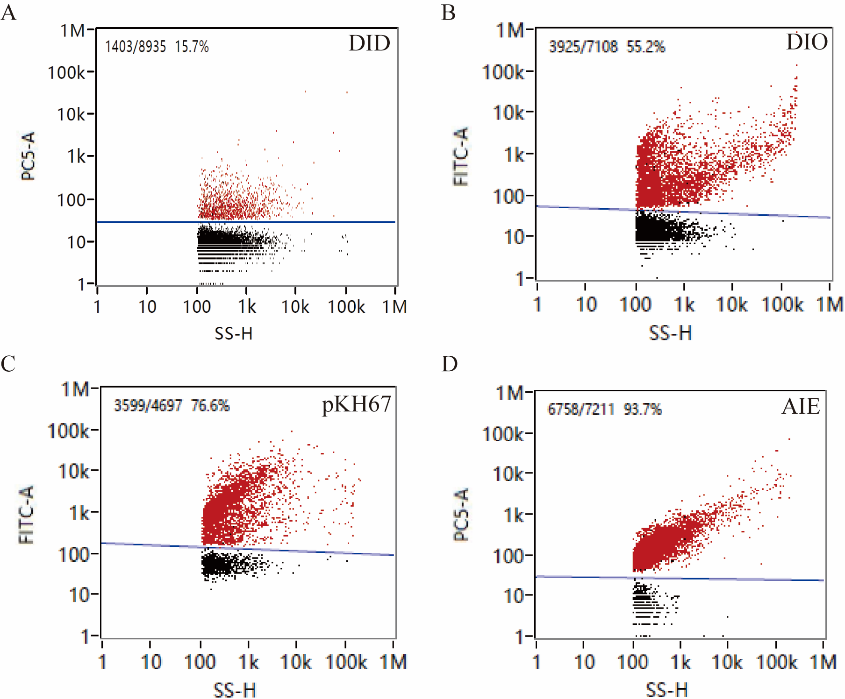

Fig. 1. (A)-(D) Fluorescent labeling of DID, DIO, pKH67 and AIE labeled MK-Exo.

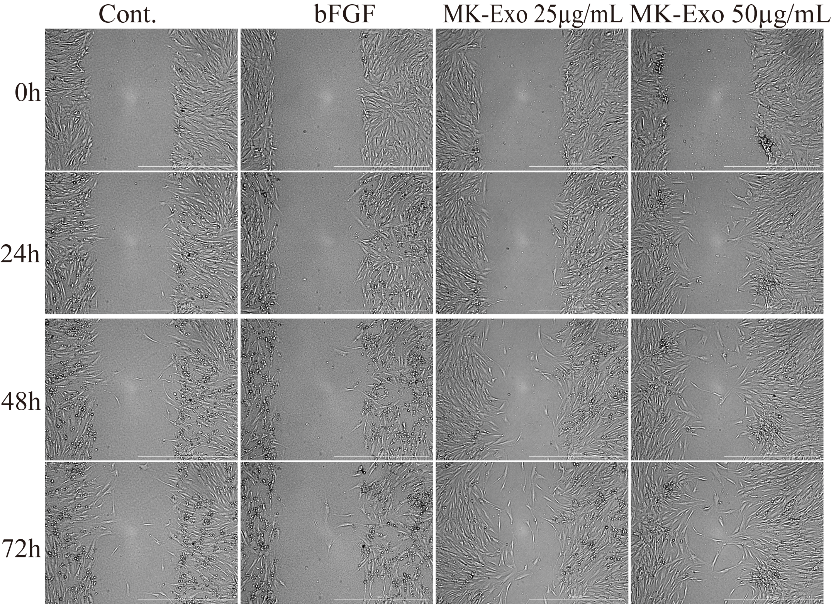

Fig. 2. Representative images of the varies of scratch width at different time points.
