## Supplementary table for "The novel roles of bovine milk-derived exosomes on skin anti-aging"

**Supplementary Tables**

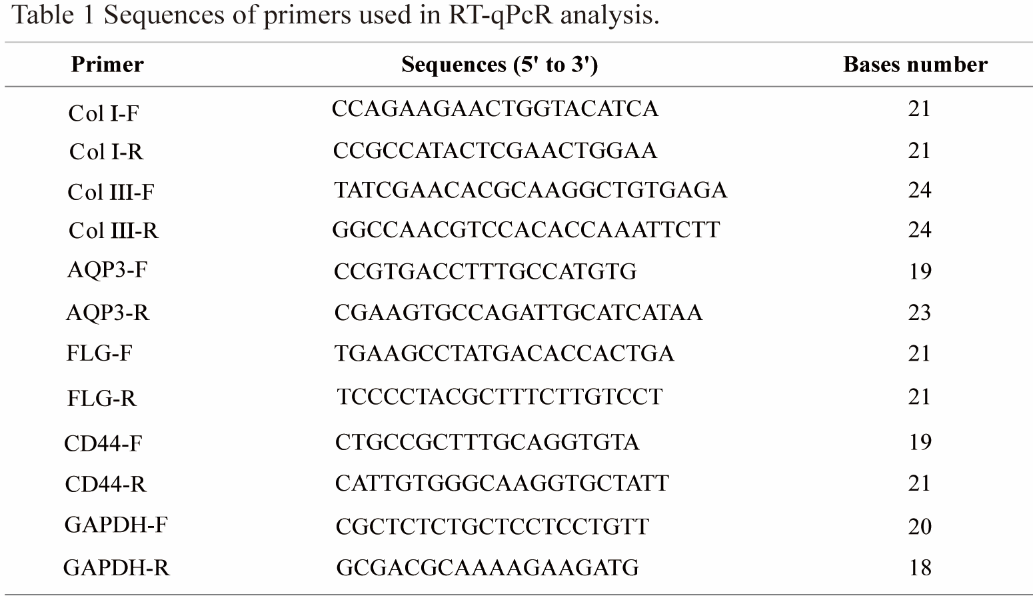

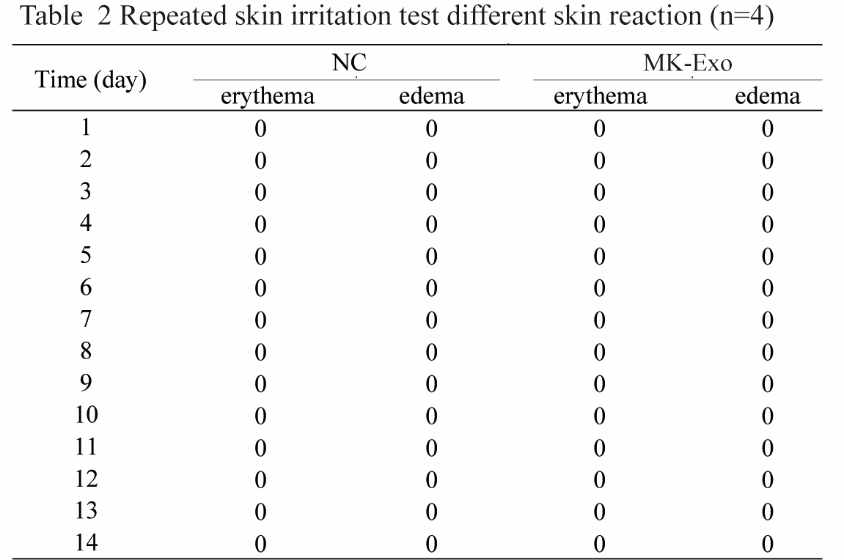

MK-Exo and the skin reaction integral were used on the New Zealand rabbits (0, 1, 2, 3, 4).

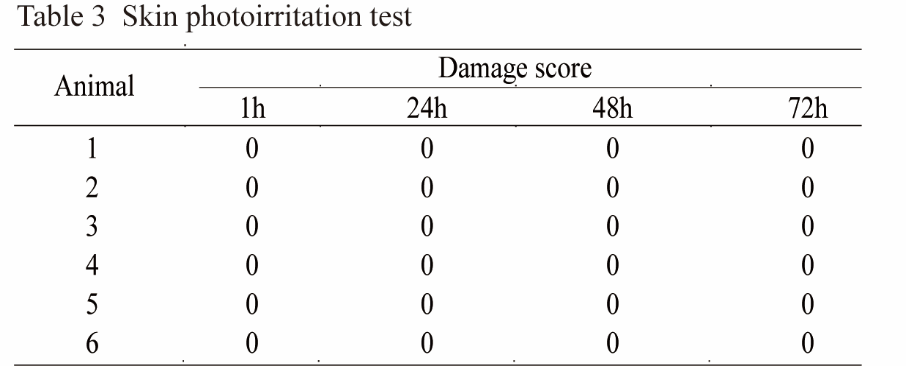

MK-Exo was used on Dunkin Hartley and determined skin reaction integral (0, 1, 2, 3, 4) (n=6).

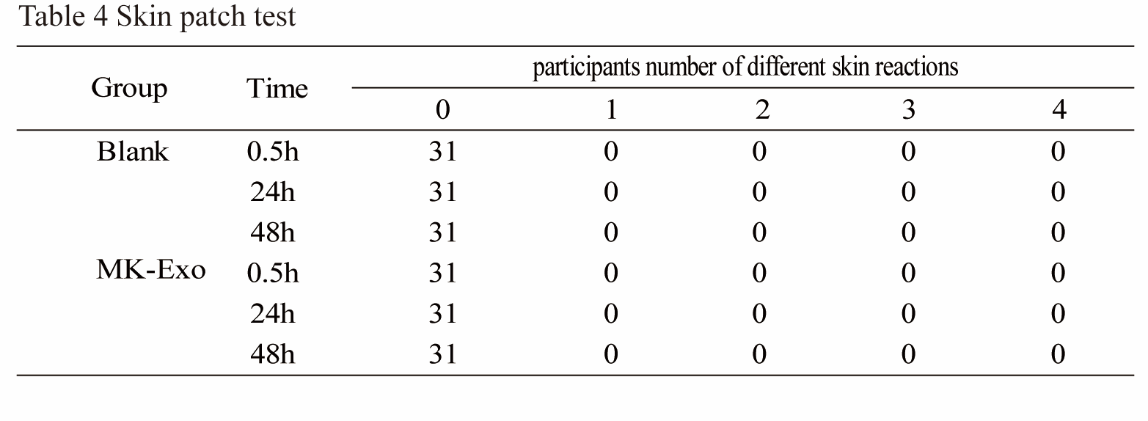

The skin reaction score (0, 1, 2, 3, 4). 0: no reaction, 1: suspicious reaction, only weak erythema, 2: weak positive reaction (erythema reaction), 3: strong positive reaction (herpetic reaction), 4: extremely positive reaction (herpetic reaction).
